# Dissecting the effects of adaptive encoding and predictive inference on a single perceptual estimation

**DOI:** 10.1101/2022.02.24.481765

**Authors:** Jongmin Moon, Oh-Sang Kwon

## Abstract

The human visual system exploits continuity in the visual environment, which induces two contrasting perceptual phenomena: repulsive aftereffects and attractive serial dependences. Recent studies have theorized that these two effects concurrently arise in perceptual processing, but empirical evidence that directly supports this hypothesis lacks. Here we show that the two effects concurrently modulate estimation behavior in a typical sequence of perceptual tasks. We first demonstrate that observers’ estimation error as a function of both the previous stimulus and response cannot be fully described by either attractive or repulsive bias but is instead well captured by a summation of repulsion from the previous stimulus and attraction toward the previous response. We then reveal that the repulsive bias is centered on the observer’s sensory encoding of the previous stimulus, which is again repelled away from the preceding trials, whereas the attractive bias is centered precisely on the previous response, which is the observer’s best prediction about the incoming stimuli. Our findings provide strong evidence that sensory coding is shaped by dynamic tuning of the system to the past stimuli, inducing repulsive aftereffects, and followed by inference incorporating the prediction from the past estimation, leading to attractive serial dependence.

## Introduction

Sensory input from the natural environment is highly structured, and humans can learn statistical regularities in sensory input and exploit them to optimize perceptual processing (Geisler, 2008; Schwartz, 2007; Simoncelli & Olshausen, 2001). Considerable advances have been made toward understanding whether and how the visual system incorporates knowledge of environmental statistics. For example, knowledge about the distribution of local orientation in natural images can be used not only to produce an efficient representation of the incoming orientation signal but also to optimize the interpretation of that orientation representation (Wei & Stocker, 2015). The relevant environmental statistics are not limited to static distributions. Most everyday visual tasks require more than processing static images, as many visual features change continuously over time (Dong & Atick, 1995; van Bergen & Jehee, 2019). For example, orientation signals arising in the natural world tend to change gradually, such that the local orientation at any given moment correlates with the orientation of the previous moment (van Bergen & Jehee, 2019). To process such dynamic sensory inputs efficiently and accurately, the visual system should consider the temporal structure between successive stimuli in the environment (Kwon et al., 2013).

There are two ways for the visual system to exploit knowledge about changing environments. First, temporal structures in sensory input provide an opportunity for the visual system to form an efficient representation of sensory information. The incoming stream of input is partially redundant because the current sensory input tends to resemble the preceding one. Therefore, the visual system can exploit the redundancy to enhance efficiency in neural coding by adaptively adjusting the response properties of sensory neurons according to the changes in input stimuli (Barlow, 1990; Brenner et al., 2000; Fairhall et al., 2001; Gepshtein et al., 2013; Gutnisky & Dragoi, 2008; Müller et al., 1999; Sharpee et al., 2006; Wainwright, 1999). The perceptual consequence of sensory adaptation is most clearly seen in repulsive aftereffects, a phenomenon in which exposure to a stimulus induces a repulsive bias in the perception of subsequent stimuli (Clifford, 2002; Clifford et al., 2007; Kohn, 2007; Thompson & Burr, 2009; Webster, 2015). Second, predictions derived from recent sensory inputs can be used to interpret the encoded signals. Numerous studies have shown that the visual system utilizes prior knowledge about the sensory environment to infer the state of the world from noisy and incomplete sensory inputs (Kersten et al., 2004; Knill & Pouget, 2004; Knill & Richards, 1996; Körding, 2007). In the context of processing dynamic sensory inputs, the current sensory input is integrated with predictions from the recent past to optimize perceptual estimation (Körding, 2007). The optimal integration model provides a normative explanation for serial dependence in perceptual behavior (Cicchini et al., 2014; Cicchini et al., 2018; Kwon & Knill, 2013; van Bergen & Jehee, 2019), a phenomenon in which perceptual estimation of the current stimulus is attracted toward the previous stimuli (Fischer & Whitney, 2014; Liberman et al., 2014).

Both repulsive aftereffects and attractive serial dependence are well-known phenomena in which the visual system leverages information from the recent past to optimize the processing of incoming visual input. These two opposite effects, however, have been historically described and theorized about in isolation. Only recently have suggestions been made that these two effects may concurrently occur in perceptual processing. The implied picture is that one effect is hidden when the other becomes behaviorally observable, but this theoretical consideration lacks direct empirical supports. Most previous studies focused on showing that either, but not both, of the two effects can be signified depending on the specifications of the experimental setup, such as stimulus features (Alais et al., 2017; Taubert et al., 2016), time interval between consecutive events (Bliss et al., 2017; Kanai & Verstraten, 2005), and task design (Czoschke et al., 2019; Fritsche et al., 2017; Pascucci et al., 2019). While these studies convergingly showed that the sequential effect varies drastically between attraction and repulsion with only a small change in experimental paradigm, they do not clarify whether and how the two effects with opposite behavioral consequences interact during the perceptual processing, which is crucial for the investigation of underlying mechanisms (Burr & Cicchini, 2014). A group of studies showed that perceptual estimates of the current stimulus are attracted to the immediately preceding stimulus and repelled away from stimuli further back in the past (or vice versa; Chopin & Mamassian, 2012; Fritsche et al., 2020; Gekas et al., 2019; Suárez-Pinilla et al., 2018). However, even these studies cannot determine whether a single sensory event can induce both attractive and repulsive biases in the immediately subsequent perception.

In this study, we test a key prediction of the current theory about attractive and repulsive sequential effects. Specifically, we reason that an observer’s current estimates would be repelled away from the previous *stimulus*, because the neural population that encodes incoming sensory information naturally adjusts its tuning characteristics according to what was encoded at the previous moment. That is, we considered the previous stimulus as a proxy for sensory measurement made by the observer in the previous trial. We also reason that, at the same time, the estimates would be attracted toward the previous *response*, because the observer’s prediction that optimizes the perceptual interpretation would be derived from what was perceived by the observer in the previous trial (Cicchini et al., 2014; van Bergen & Jehee, 2019). In the absence of feedback, the observer does not have direct access to the true stimulus in the previous trial, so the final perception in the previous trial is the best estimate the observer has about the previous stimulus. Thus, although it may appear trivial at first glance to determine whether it is a stimulus and/or response that biases the subsequent perception, it has important implications on how and why the visual system processes sensory information according to the recent sensory events.

To test this hypothesis, we aimed to estimate the concurrent influence of the previous stimulus and response on current perception. In a series of trials, we asked subjects to view a field of moving dots and to report the perceived direction of motion. The subjects’ responses to the direction of motion were systematically attracted toward the direction of motion in the previous trial, consistent with previous studies. Importantly, however, by representing the response error as a function of both the previous stimulus and response, we demonstrated that the estimation responses are repelled from the direction of motion in the previous trial, resembling the repulsive aftereffects, and at the same time attracted toward the reported direction of motion in the previous trial, mirroring the attractive serial dependence.

## Results

Processing a sequence of images involves interactions between the current input and past observations, and the repulsive aftereffect and the attractive serial dependence are regarded as behavioral consequences of these interactions (Burr & Cicchini, 2014; Thompson & Burr, 2009). We sought to investigate the determinants of these interactions by separately estimating the respective effects of stimulus and response on subsequent perception. In a typical perceptual estimation task, however, the value of a given stimulus and the subjects’ response to it would be strongly correlated, making it challenging to distinguish which of those two is related to the observed bias. To effectively disentangle them, we carefully modulated the stimulus dynamics across subjects, thereby ensuring the collection of a sufficient number of informative trials. Subjects viewed a random-dot motion stimulus whose direction of motion randomly varied from the direction of motion in the previous trial following a uniform distribution (range: ±20°, 40°, 80°, or 180°), and reported the perceived direction of motion (**Fig. 1a**).

**Figure 1.**
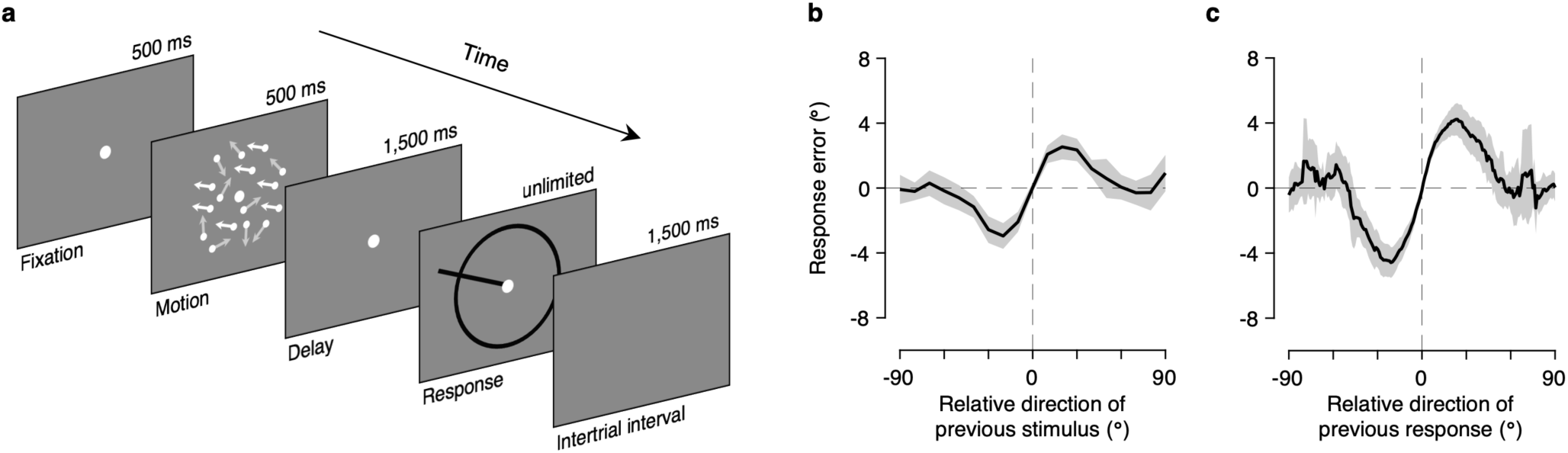
Task procedure and marginal bias plots. **(a)** On each trial, subjects viewed a random-dot motion stimulus and reported the perceived direction of motion by extending a dark bar from the center point. They were instructed to report the direction of motion of dots by swiping their finger on a touchpad to extend a bar from the center of the display to the direction of their estimate and confirm that report with a click on the touchpad. **(b)** Bias plot marginalized on the relative direction of the previous response. Response errors are expressed as a function of the relative direction of the previous stimulus (i.e., previous stimulus minus current stimulus). Consistent with earlier studies, estimation responses are systematically biased to the previous stimulus. **(c)** Bias plot marginalized on the relative direction of the previous stimulus. Response errors are expressed as a function of the relative direction of the previous response (i.e., previous response minus current stimulus). Again, estimation responses are systematically biased to the previous response. Shaded regions represent 95% confidence intervals.

### Attractive sequential effects in motion direction estimation

We first examined whether we successfully replicated previously reported serial dependence. To do this, we adopted the standard approach from earlier studies that focused on the effect of previous stimuli on current perception (Fischer & Whitney, 2014; Liberman et al., 2014). First, the response error was computed as the angular difference between the reported and presented motion directions, with positive angles corresponding to clockwise deviations. In doing this, we corrected for the well-known estimation biases toward or away from the cardinal directions (Kosovicheva & Whitney, 2017; Wei & Stocker, 2015), which could have been confounded with the sequential effects (see **Methods**). We then examined the patterns of subjects’ response errors as a function of the relative direction of the previous stimulus to the current stimulus (previous stimulus direction minus current stimulus direction). As expected, we found that the estimation responses were systematically biased to stimuli that were previously viewed (**Fig. 1b**). We also examined the response error as a function of the relative direction of the previous response to the current stimulus (previous response direction minus current stimulus direction) and found that the responses were again attracted toward responses that were recently made (**Fig. 1c**). These sequential effects were highly significant (previous stimulus: *p* < 0.001; previous response: *p* < 0.001, hierarchical Bayesian parameter estimation; see **Methods**), and their magnitudes and attraction profiles were consistent with earlier reports (Fischer & Whitney, 2014; Fritsche et al., 2017; Pascucci et al., 2019), following a first derivative of Gaussian (DoG) curve (previous stimulus: amplitude = 2.93°, peak location = 20.87°; previous response: amplitude = 4.62°, peak location = 21.02°).

However, plotting response errors as a function of either previous stimulus or response direction does not reflect the isolated effect of the previous stimulus or response, since the direction of motion stimulus and the subjects’ corresponding response to it are strongly correlated: The Pearson’s correlation between the true and reported direction of motion had a median (and interquartile range) of 0.910 (0.884–0.929) across subjects. Therefore, it would be inappropriate to judge which component of the previous trial truly influenced the response of the current trial based only on the patterns of biases marginalized over the previous stimulus or response (**Fig. 1b,c**).

### Disentangling the effects of previous stimuli and responses

The subjects’ estimation behavior in the current trial could have been influenced by the direction of the preceding stimulus, subjects’ perceptual estimate of it, or both. To obtain a general insight into which aspects of the previous trial affected subjects’ performance and how, we visualized subjects’ response error in a two-dimensional map as a function of both previous stimulus and response directions (**Fig. 2a**; see **Supplementary Figure 1** for individual subjects’ data). In this joint bias map, the x-axis represents the relative direction of the previous stimulus, the y-axis represents the relative direction of the previous response, and the color of each pixel represents the subjects’ response error. Pixels with warm colors indicate positive errors, and pixels with cool colors indicate negative errors. Thus, both warm-colored pixels with positive labels on the axis and cool-colored pixels with negative labels on the axis represent attractive biases, whereas warm-colored pixels with negative labels on the axis and cool-colored pixels with positive labels on the axis represent repulsive biases.

**Figure 2.**
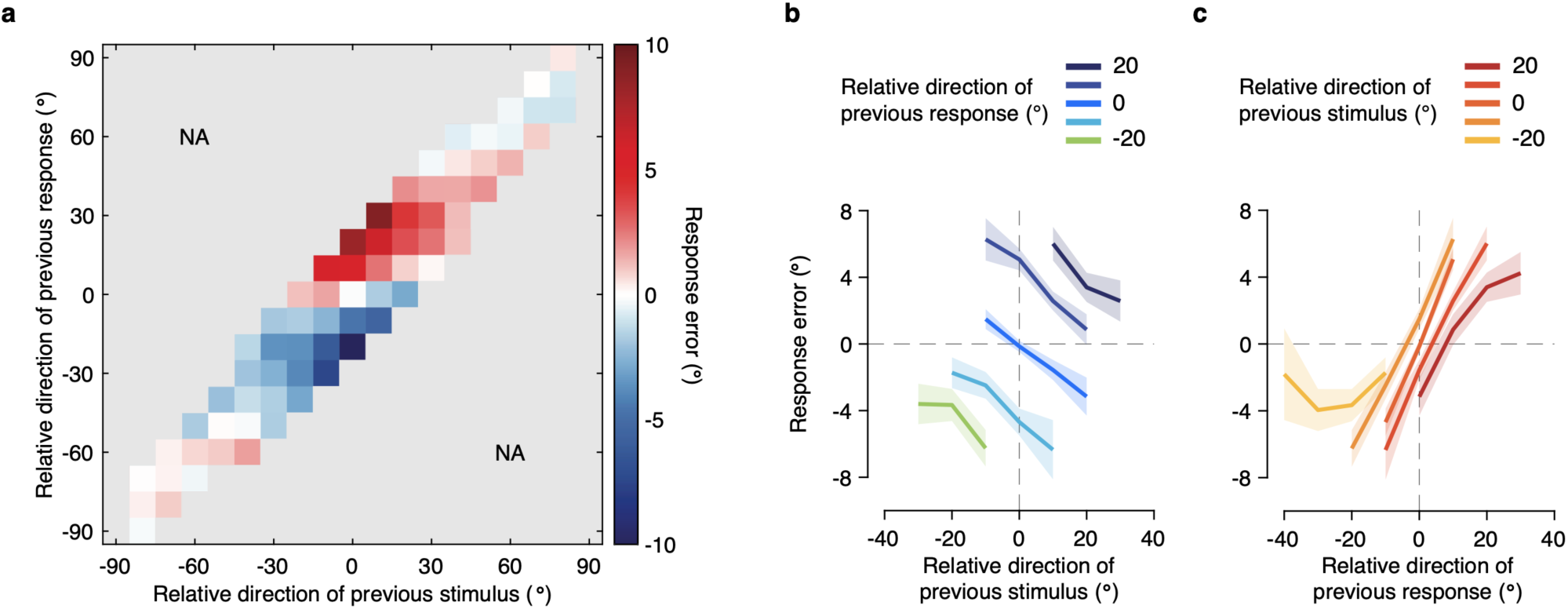
Joint bias map and conditional bias plots. **(a)** Joint bias map. Response errors are plotted as a function of both the previous stimulus and response direction relative to the current stimulus. For each subject, we binned the response errors within 10° bins according to the relative direction of the previous stimulus and response. Group means with less than sixteen subjects’ data were excluded before illustration. The color of each pixel represents response error on the current trial. Positive values on the x- and y-axis indicate that the previous stimulus and response direction was more clockwise than the current stimulus, respectively, and positive errors indicate that the estimated direction was more clockwise than the true stimulus direction. Estimation responses on the current trial are systematically repelled away from the stimulus direction of the previous trial, while they are strongly attracted toward the response direction of the previous trial. NA, not available. **(b)** Bias plot conditioned on relative direction of previous response. Response errors split by relative direction of the previous response were plotted as a function of relative direction of the previous stimulus. Bins with less than five trials within a subject and group means with less than sixteen subjects’ data were excluded before illustration. As can be seen across five lines with negative slopes, estimation responses are negatively biased away from the previous stimulus. **(c)** Bias plot conditioned on relative direction of previous stimulus. Response errors split by relative direction of the previous stimulus were plotted as a function of relative direction of the previous response. Estimation responses are positively biased toward the previous response, as indicated by the positive slopes. Shaded regions represent 95% confidence intervals.

Visual inspection of the joint bias map reveals that it has a rich structure beyond the traditional view of the sequential effect. Specifically, it becomes immediately apparent from the map that the sequential effect cannot be fully characterized by a simple attractive or repulsive bias and is not solely determined by either the previous stimulus or the previous response. Instead, errors in direction estimates seem to be driven by both attractive and repulsive biases that depend systematically on both the previous stimulus and previous response, relative to the current stimulus. In the following, we further examine the characteristic features of the joint bias map one by one.

Our first and most crucial observation was that the estimation responses were systematically repelled away from the stimulus direction of the previous trial, while they were strongly attracted toward the response direction of the previous trial. Consider, for example, horizontally neighboring pixels labeled as zero on the y-axis. Previous responses relative to the current stimulus (i.e., y-axis value) are fixed across these pixels, so they would reveal the sole effect of the previous stimulus. These pixels show that the response error decreases as the relative stimulus direction of the previous trial increases, suggesting that the effect of the previous stimulus is repulsive. This indicates that when the random dots in the previous trial moved in a direction more clockwise than the direction of motion of the dots in the present trial, subjects perceived the present random dots as moving in the direction more counterclockwise than their true direction of motion. The repulsive effect of the previous stimulus becomes even more evident when we plot the response error, split by the relative direction of the previous response, as a function of the relative direction of the previous stimulus (**Fig. 2b**). This would reveal the sole effect of the previous stimulus on the current response, since the effect of the previous response is fixed within each line. The bias plot clearly shows negative slopes regardless of previous response directions, which confirms the robust repulsion away from the previous stimulus direction.

In contrast, vertically neighboring pixels that are labeled zero on the x-axis show that the response error increases as the relative response direction of the previous trial increases, suggesting that the effect of the previous response is attractive. This indicates that when subjects perceived and subsequently reported that the random dots in the previous trial had moved in the direction more clockwise than the direction of motion of dots in the present trial, they perceived the present random dots as moving in the direction more clockwise than their true direction of motion. Again, the attractive effect of the previous response becomes even more pronounced by plotting the response error split by the relative direction of the previous stimulus as a function of the relative direction of the previous response (**Fig. 2c**). The bias plot exhibited positive slopes, confirming the strong attraction toward the previous response direction.

The joint bias map (**Fig. 2a**) also provides us with an idea about how these two opposite effects interact. Based on the linearly increasing response errors from the lower right to the upper left, we can observe that these two sequential biases are combined in an additive manner. Consequently, the bias was strongest when the directions of the previous stimulus and response were opposite relative to the current stimulus, as revealed by the response errors on the second and fourth quadrants, with their magnitudes being up to 10° on average.

Lastly, we confirmed that it would be inappropriate to quantify the biases with a linear regression that uses the relative direction of the previous stimulus and response as predictors. Both repulsion away from the previous stimulus and attraction toward the previous response were observed when the previous stimulus and response values were similar to the current stimulus, but these patterns were diminished or even slightly reversed when they were very different (**Fig. 2a**), suggesting that linear regression would not be an appropriate way to quantify the biases.

### Additivity of concurrent attractive and repulsive sequential effects

We used a descriptive model to simultaneously quantify the attractive and repulsive biases. Based on existing literature, we assumed that patterns of the attractive and repulsive effects would each resemble the DoG curve and that the two effects would linearly add up to compose the resulting sequential biases in subjects’ estimation behavior. We fitted a sum of two independent DoG curves, each representing the bias to or away from the previous stimulus and bias to or away from the previous response, respectively. In the following, we refer to this model as the Stimulus & Response model, as both the previous stimulus and previous response bias the current perception. We compare the Stimulus & Response model to alternative models in which either previous stimulus or previous response biases the current estimates (designated as the Stimulus model and the Response model, respectively). We first predicted the estimation behavior using the best-fitting parameters of each model and plotted joint bias maps to examine the model performance (**Fig. 3a-c**). While the behaviors of neither Stimulus model (**Fig. 3a**) nor Response model (**Fig. 3b**) properly emulated the human data (**Fig. 2a**), the Stimulus & Response model successfully predicted the pattern of biases observed in the empirical data (**Fig. 3c**). Models were also quantitatively compared using the Akaike information criterion (AIC) to account for differences in model complexity, with data reported as the mean AIC difference from the best-fitting model followed by a 95% confidence interval in brackets. The Stimulus & Response model outperformed the Stimulus model (AIC difference = 143.4 [74.5 212.3]) and the Response model (AIC difference = 39.5 [24.5 54.4]; **Fig. 3d**).

**Figure 3.**
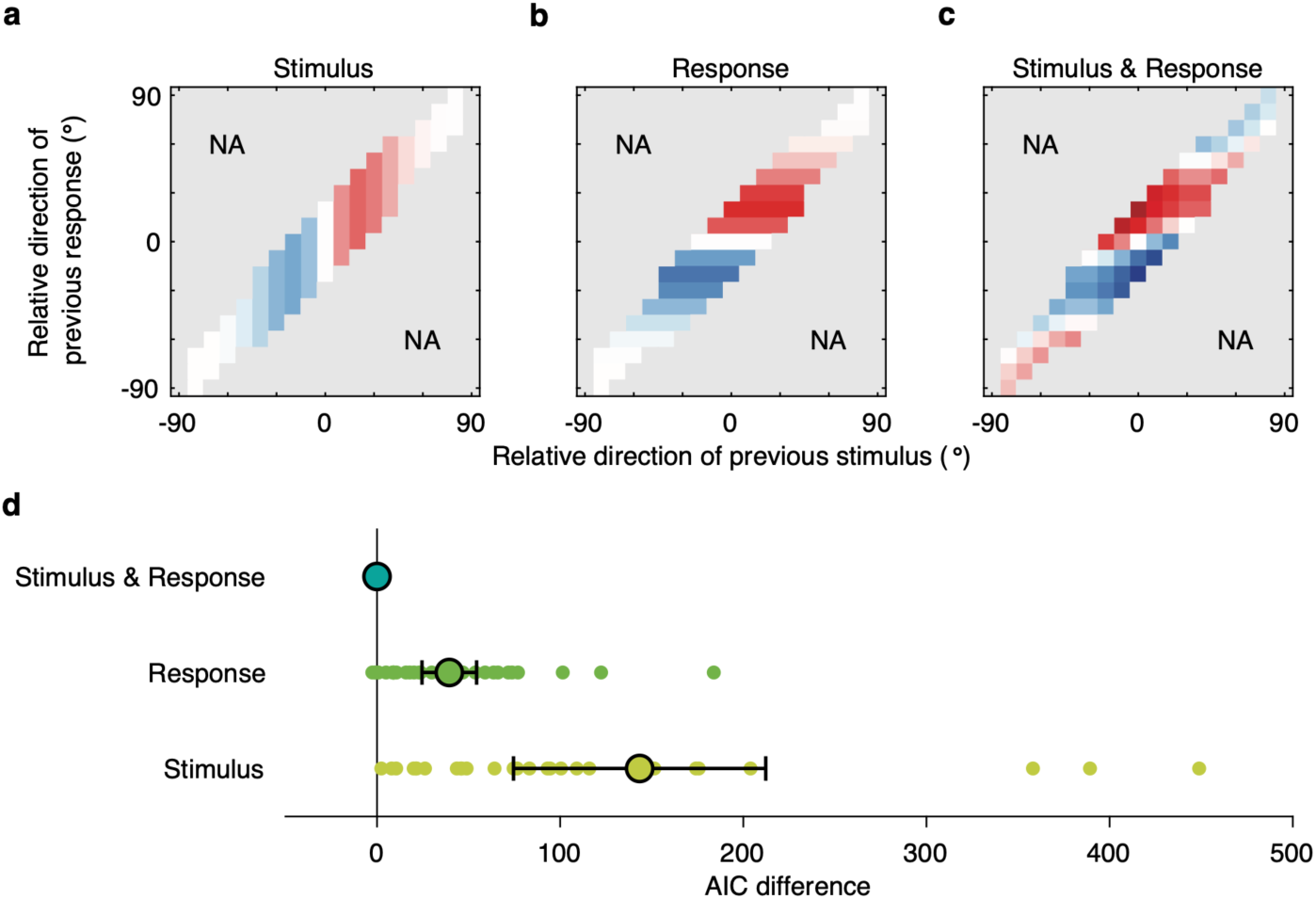
Model comparison. **(a)** Prediction from a single DoG function of relative direction of the previous stimulus (Stimulus model). Color conventions are as described in **Figure 2a**, and pixels are shown for only those available in **Figure 2a. (b)** Prediction from a single DoG function of relative direction of the previous response (Response model). **(c)** Prediction from the linear sum of two independent DoG functions, each representing bias to (or away from) the previous stimulus and bias to (or away from) the previous response (Stimulus & Response model). Characteristic patterns of the joint bias map are in excellent agreement with empirical data shown in **Figure 2a. (d)** Model comparison using Akaike information criterion (AIC). Dots represent individual subject differences in AIC values between each model and the Stimulus & Response model. Circles and error bars represent mean and 95% confidence intervals. Positive numbers represent a worse fit than the Stimulus & Response model. The Stimulus & Response model fits the best.

We did not constrain the direction of the biases in these models to be attractive or repulsive. Nevertheless, the fitting results of the Stimulus & Response model revealed that the previous stimulus repelled the subsequent responses, while the previous response attracted them (previous stimulus: amplitude = −4.97°, *p* < 0.001, previous response: amplitude = 8.52°, *p* < 0.001; **Fig. 4a**), consistent with our visual inspection of the joint bias map. Note that the absolute magnitudes of both attractive and repulsive biases are considerably larger than those observed in the marginal bias plots (**Fig. 1b,c**), as well as the typical sequential biases reported in the literature. The attraction toward the previous response was significantly larger in absolute magnitude than the repulsion away from the previous stimulus (*p* < 0.001), accounting for the net attractive bias observed in the marginal bias plots of our data (**Fig. 1b,c**) and in earlier studies. In addition, the repulsion away from the previous stimulus was more broadly tuned than attraction toward the previous response (previous stimulus: peak location = 34.89°; previous response: peak location = 27.10°; difference: *p* < 0.001; **Fig. 4b**), which is consistent with the small repulsive bias often observed in earlier studies when successive stimuli were markedly different (Fritsche et al., 2017; Bliss et al. 2017; Samaha et al., 2019; Fritsche & de Lange, 2019). Overall, our results show that the observed patterns of sequential effects in the joint bias map can be well described by the linear sum of two curves characterizing the attraction toward the previous response and repulsion away from the previous stimulus.

**Figure 4.**
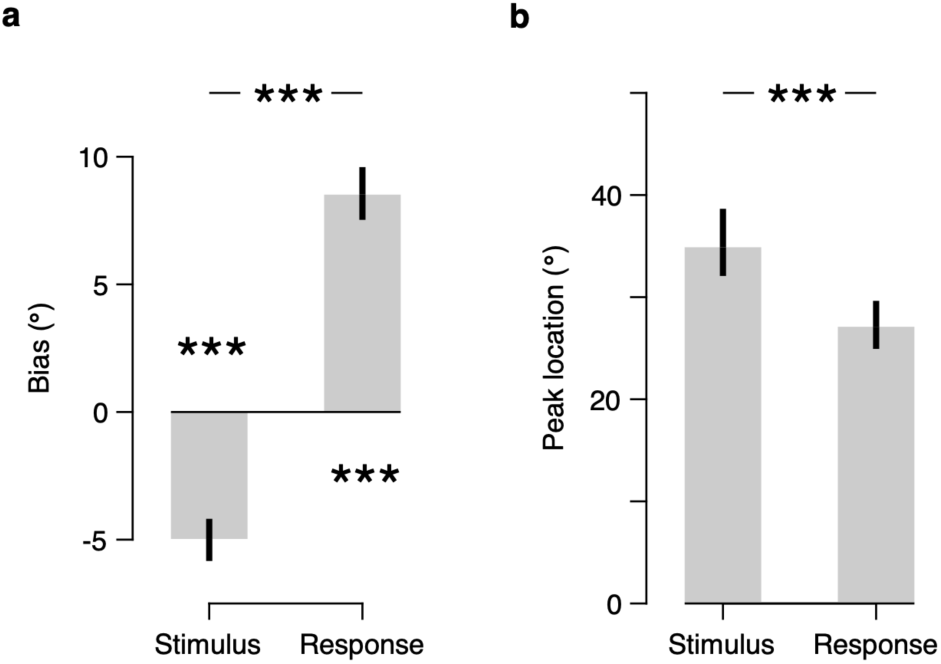
Quantifying the two opposite biases. **(a)** Estimated bias magnitudes in the Stimulus & Response model. Bias is negative for the previous stimulus and positive for the previous response, meaning that the estimation responses were repelled away from the previous stimulus and attracted toward the previous response. Both biases are highly significant, with the bias to the previous response being significantly larger in absolute magnitude than the bias away from the previous stimulus. **(b)** Estimated peak locations of the bias curves in the Stimulus & Response model. Bias away from the previous stimulus is more broadly tuned than the bias to the previous response. Error bars represent 95% credible intervals. \*\*\**p* < 0.001, hierarchical Bayesian parameter estimation.

### Estimating the centers of biases

We constrained the Stimulus & Response model in such a way that the centers of the two DoG curves for repulsive and attractive biases are located at the previous stimulus or response. The rationale was that the previous stimulus is a proxy for the tuning state of the neural population in the encoding stage, which may induce repulsion, and the previous response is a proxy for the prediction from the past estimation, which may induce attraction. However, it is an open question as to whether the previous stimulus and response are indeed the centers of bias. For instance, it is plausible that the tuning characteristics of the neural population are affected by not only the previous stimulus but also the stimulus before the previous stimulus, while the prediction from the past estimation is indeed centered at the previous response that reflects the observer’s best estimate of the previous stimulus, equivalent to the best predictor for the incoming input in our task.

To estimate the centers of biases, we fitted a new model in which the centers of the attraction and repulsion curves were parameterized as *β*Response_*n*−1_ + (1 − *β*)Stimulus_*n*−1_. We refer to this model as the Attraction & Repulsion model. Note that whereas the Stimulus & Response model predetermines the centers of bias curves, the Attraction & Repulsion model constrains the direction of bias curves (i.e., one bias curve is for attraction and the other is for repulsion). If *β* is one, the current response is biased to (or away from) the previous response, and if *β* is zero, the current response is biased to (or away from) the previous stimulus. We fitted the model to data as for the Stimulus & Response model, but with two *β*s, each for attraction (*β*_*att*_) and repulsion (*β*_*rep*_) curves, as additional free parameters. For the attraction curve, we found that *β*_*att*_ was significantly larger than zero (*p* < 0.001) and tightly clustered around one (**Fig. 5a**, red circle). This provides additional support for the attraction toward the previous response and not toward the previous stimulus (or a combination of the two). We also found that, for the repulsion curve, *β*_*rep*_ is vastly different from one (*p* < 0.001; **Fig. 5a**, blue circle) and closer to zero, suggesting that the current response is not repelled from the previous response. However, the estimated *β*_*rep*_ was not exactly clustered around zero (*p* < 0.001). Instead, *β*_*rep*_ was slightly but consistently negative across subjects, indicating that the center of the repulsion curve was located near the previous stimulus but slightly shifted away from the previous response.

**Figure 5.**
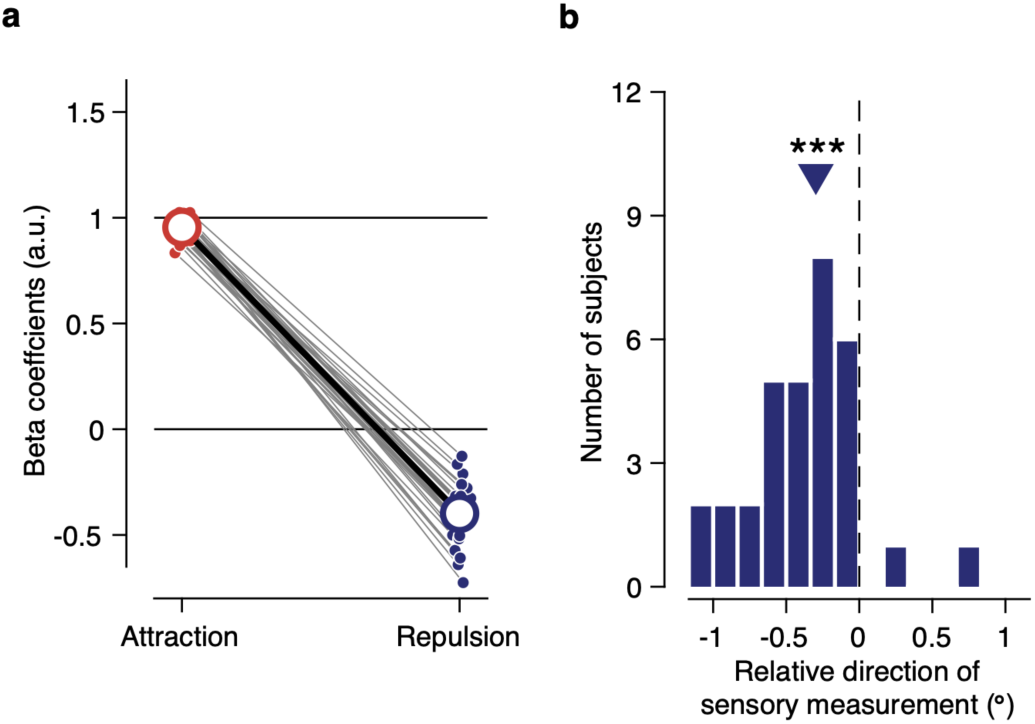
Estimating the centers of biases. **(a)** Estimated center of the attraction and repulsion biases in the Attraction & Repulsion model. Dots represent the best-fitting beta coefficients for individual subjects, and open circles represent the population mean. Beta coefficient zero indicates that the bias is toward (or away from) the previous stimulus, and one indicates that the bias is toward (or away from) the previous response. Center of attraction is located on the previous response, while center of repulsion is located near the previous stimulus but slightly shifted away from the response. **(b)** Overall shift in sensory measurement away from the response. Sensory measurements were first computed by repelling the current stimulus away from the previous stimulus using the best-fitting parameters of the repulsive bias in the Stimulus & Response model. They were then flipped depending on the sign of response error on each trial such that positive numbers represent a shift in sensory measurement toward the response, and negative numbers represent a shift away from the response, relative to the stimulus. The triangle represents the median of the distribution. Sensory measurements were on average shifted away from the response, relative to the stimulus, consistent with the estimated center of repulsion in **a**. \*\*\**p* < 0.001, Wilcoxon signed-rank test.

We propose that the systematic deviation of *β*_*rep*_ from the previous stimulus indicates that the center of repulsion is determined by “sensory measurement” of the previous stimulus rather than the previous stimulus itself. The sensory measurement is defined as the latent value that represents the measured sensory input, which is affected by the repulsive bias in encoding but has not yet been affected by the attractive bias in decoding. Consider a case in which the current response is located clockwise relative to the current stimulus. It is likely that the previous stimulus is also located clockwise relative to the current stimulus because the response is biased, on average, to the previous stimulus. Consequently, the sensory measurement of the current stimulus would systematically deviate from the current stimulus in a direction away from the current response. We ran a simulation using the best-fitting parameters of the repulsive bias in the Stimulus & Response model to examine the mean deviation of the sensory measurement from the stimulus. The results showed an overall shift in sensory measurement in a direction away from the response (median = −0.30°; interquartile range = 0.43°; *p* < 0.001, Wilcoxon signed-rank test; **Fig. 5b**). In short, if the sensory measurement determines the center of repulsion, it is expected that the center of repulsion deviates from the previous stimulus in a direction away from the previous response (**Fig. 5b)**, which is exactly what we observed in our data (**Fig. 5a**).

## Discussion

Our natural environment is abundant with statistical regularities, a typical one being that the environment is stable over time (Dong & Atick, 1995; van Bergen & Jehee, 2019). If it is scorching hot today, no one would expect heavy snow the next day. It is no surprise that an observer utilizes such environmental stability to improve their perceptual performance. However, there is a certain irony in this situation because an ideal observer should not only ensure that sensory neurons adapt to the changes in input stimuli, repelling the current perception, but also perform probabilistic inference based on predictions from past observations, attracting the current perception toward the recent history. These two have been widely suggested as underlying mechanisms of the repulsive aftereffects and attractive serial dependence, but it remains unclear whether and how these two opposite effects interact in the perceptual process (Burr & Cicchini, 2014). Here, we studied the concurrent attractive and repulsive sequential effects in visual perception to determine how these competing demands are resolved during visual processing. Specifically, we showed that the previous stimulus repels the current response, the previous response attracts the current response, and these two biases add up to shape a unique pattern in estimation behavior. We also directly estimated the centers of the biases and confirmed that whereas the attraction bias is centered precisely at the previous response, the repulsion bias is centered at the sensory measurement of the previous stimulus. We believe that the concurrent attractive and repulsive sequential effects could be present in many other data in literature (see also Sadil et al., 2021), as the overall attraction bias found in our study mirrors several previous reports showing similar attractive biases in numerous perceptual and cognitive domains.

### Concurrency and additivity of attractive and repulsive biases

The current results further develop our understanding of sequential effects in two main ways. First, this study dissociated the attractive and repulsive effects with the same task throughout the experiment and without further manipulation in experimental procedures and provided a clear picture of the concurrency of the two opposite biases. Although suggestions have been made on the possibility that the two opposite biases coexist (Fritsche et al., 2017; Pascucci et al., 2019; Sadil et al., 2021), their nature and origin remain topics of debate, partly due to a lack of direct empirical support. Notably, if repulsion away from the previous stimulus and attraction toward the previous response occur at the same time, then the bias would be exceedingly strong when the directions of the previous stimulus and response are opposite relative to the current stimulus. This is exactly what we found (**Fig. 2a**). Second, we empirically demonstrated that the effects of stimulus and response of the preceding trial are largely additive in the perception of the current trial. We fitted the linear sum of two curves representing biases to (or away from) the previous stimulus and response, respectively, and found that it could capture the unique pattern in the subjects’ sequential estimation (**Fig. 4c**). These results suggest that the two contrasting biases originate from independent mechanisms and open new directions for research on sequential effects.

### Rapid adaptation to visual stimulus

Our results support the idea that the visual system adapts to changes in briefly presented stimuli, resulting in a repulsive bias away from the previous stimulus. Sensory adaptation and repulsive aftereffects have historically been studied in the context of prolonged stimulus exposure, but increasing evidence also suggests a more rapid form of adaptation. Such adaptation is known to occur rapidly following a stimulus exposure as brief as a few hundred, or even tens, of milliseconds (Dragoi et al., 2002; Fairhall et al., 2001; Glasser et al., 2011; Gutnisky & Dragoi, 2008; Müller et al., 1999) and persist over dozens of seconds despite the presentation of several intervening stimuli (Fritsche et al., 2021), inducing robust repulsive biases in subsequent behavioral reports (Aagten-Murphy & Burr, 2016; Alais et al., 2017; Bliss et al., 2017; Chopin & Mamassian, 2012; Czoschke et al., 2019; Fritsche et al., 2017; Fornaciai & Park, 2019; Glasser et al., 2011; Kanai & Verstraten, 2005; Pascucci et al., 2019; Taubert et al., 2016) over a prolonged timescale (Fritsche et al., 2020; Gekas et al., 2019; Suárez-Pinilla et al., 2018). A recent neuroimaging study also found that sensory representations in the visual cortex were significantly and substantially repelled from the previous stimulus even when stimuli were presented only for a second, and successive stimuli were separated by at least ten seconds (Sheehan & Serences, 2021). Our findings are consistent with all these documentations, which together confirm that the visual system translates input stimuli into sensory representations in an efficient way, inducing a repulsive bias in subsequent perception, even when the stimuli are only briefly presented.

### Characteristics of attraction bias

The attractive bias toward the previous response does not necessarily imply that subjects merely reproduced their previous motor commands. Several studies have experimentally precluded the effect of motor replication and observed an attractive bias even when reproducing the previous motor command would not result in such attraction (Cicchini et al., 2017; Fischer & Whitney, 2014; Kowler, 1989; Kwon & Knill, 2013; Murai & Whitney, 2021). For example, when subjects were asked to make a flipped version of orientation response that is vertically symmetric to the stimulus orientation on every second trial, they still systematically biased their responses toward the previous stimulus, and not toward the previous motor response, suggesting that the bias was introduced during the perceptual process (Cicchini et al., 2017). Recent neurophysiological studies also denied the effect of motor replication by showing that the biases are already present in neural representations in the visual cortex (John-Saaltink et al., 2016) even without an explicit task (Fornaciai & Park, 2018). Therefore, care must be taken when interpreting the attractive biases to the previous responses reported in this study. Our proposal is that the previous response, to which the observer’s current estimation responses are attracted, is the observer’s best estimate of the stimulus in the previous trial. Our sensors can only provide noisy measurements of a given stimulus, so the visual system does not have direct access to the true stimulus value. Instead, the system infers the stimulus value from the sensory measurement and creates a perceptual estimate that sometimes deviates on average from the true stimulus value, as in our experiment. Because it is the final estimate from which the visual system predicts the subsequent stimulus, it is natural for the system to bias its current estimate toward the previous estimate, rather than toward the previous stimulus (Cicchini et al., 2014; van Bergen & Jehee, 2019).

### Computational basis of concurrent repulsion and attraction biases

We now consider a generative computational model to confirm that the proposed conceptual framework can be instantiated in a plausible encoding-decoding system. We assume that direction estimates are the outcome of a perceptual encoding-decoding process, beginning with sensory input to a population of direction-selective neurons with independent Poisson spike count variability. The sensory input and uncertainty associated with it are naturally encoded in the population activity of these neurons in the form of the likelihood function (Jazayeri & Movshon, 2006; Li et al., 2021; Ma et al., 2006; Walker et al., 2020). The encoding stage of the model was further characterized by the adaptation of these neurons to the direction of the preceding motion. Motivated by a vast amount of neurophysiological evidence (Clifford, 2002; Clifford et al., 2007; Kohn, 2007; Webster, 2015), we assume that the primary effect of sensory adaptation is a reduction in the response gain of neurons selective to the direction of preceding motion (**Fig. 6a**). The simulation of this encoder model confirms that such changes in the neural coding result in repulsive shifts of the average likelihood function away from the preceding motion direction (**Fig. 6b**). Due to sensory noise and adaptation, the sensory measurement of the preceding motion direction may differ from the true direction of the preceding motion (**Fig. 5**). To this end, we also considered an alternative scenario in which the neurons adapt to the sensory measurement (i.e., gain reduction of a neuron according to its spike count in the previous trial), rather than the true stimulus value, and confirmed that the likelihood function is still, on average, repelled away from the preceding motion direction. It is worth noting that adaptation schemes need not be restricted to gain reduction but can be easily generalized to other changes in the tuning mechanism, such as shifts in the neuron’s preferred direction (Dragoi et al., 2002; Müller et al., 1999) or flank suppression (Kohn & Movshon, 2004), which are known to generate similar repulsive effects (see also Seriès et al., 2009; Stocker & Simoncelli, 2006).

**Figure 6.**
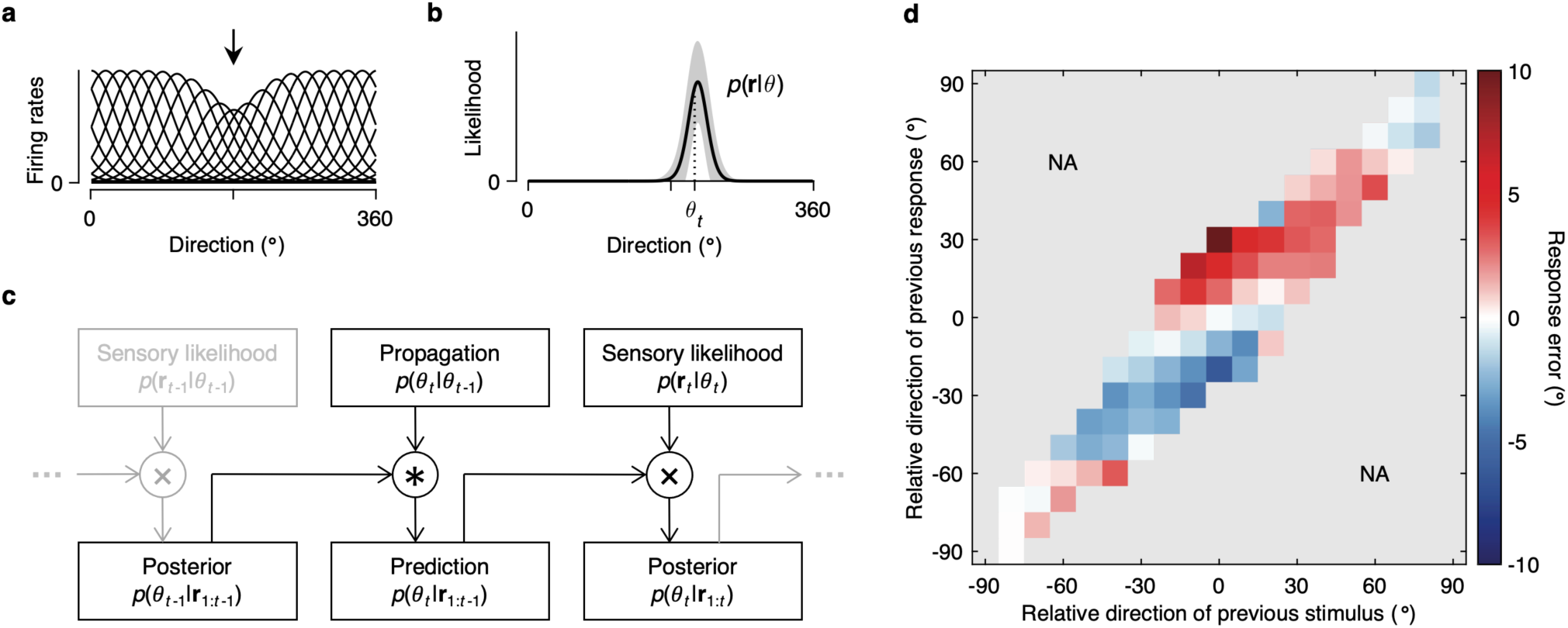
Generative model. **(a)** Tuning curves after adaptation at 180° (black arrow). Adaptation was modeled as a reduction of the response gain in neurons selective to the adapter direction. **(b)** Population likelihood. Likelihoods were computed assuming independent Poisson spike count variability and averaged over 1,000 presentations of the stimulus direction *θ*_*t*_, each following a presentation of the adapter direction as in **a**. Due to adaptation, the likelihoods are on average shifted away from the adapter direction. Shaded regions represent ±1 s.d. **(c)** Recurrent Bayesian inference. The model first convolves posterior belief on the previous stimulus *p*(*θ*_*t*−1_|**r**_1: *t*−1_) with the propagation noise distribution *p*(*θ*_*t*_|*θ*_*t*−1_), to predict the current stimulus *θ*_*t*_ based on the past sensory responses **r**_1: *t*−1_. The model then combines this prediction *p*(*θ*_*t*_|**r**_1:*t*−1_) with the new sensory information *p*(**r**_*t*_|*θ*_*t*_), to produce posterior belief on the current stimulus *p*(*θ*_*t*_|**r**_1:*t*_). **(d)** Model simulation. Simulation data were analyzed and plotted as in **Figure 2a**. The model successfully generated the characteristic features of the human estimation behavior.

Next, we assumed that the model observer uses knowledge about the dynamics of the stimulus to convert its belief about the stimulus value in the previous trial into a prediction about the stimulus value in the subsequent trial (Körding, 2007; van Bergen & Kriegeskorte, 2020). Motivated by previous work (van Bergen & Jehee, 2019), we modeled knowledge about the changes in stimuli within a given time interval (i.e., propagation noise) as a mixture of uniform and von Mises distributions with zero mean. The model observer convolved this distribution with posterior belief on the previous stimulus to make predictions about the current stimulus, which is then combined with the new sensory information to produce a posterior belief on the current stimulus based on the past and current sensory information (**Fig. 6c**). This computation follows a long history of research that models perceptual and sensorimotor behavior as recursive Bayesian inference (Körding et al., 2007; Kwon et al., 2015; Mehta & Schaal, 2002; Orban de Xivry et al., 2013; Saunders & Knill, 2004; Wolpert et al., 1995). We simulated the model observer’s estimation responses to the sequence of stimuli that our subjects encountered, assuming a squared-error loss function and no motor noise. The results showed that our model observer produced a joint bias map (**Fig. 6d**) that was close to that of the human subjects (**Fig. 2a**). The estimation responses of the model observer were repelled away from the previous stimulus because adaptation shifted the likelihood function and attracted toward the previous response because the prediction was made from the observer’s belief in the previous trial. The fact that this observer model can approximate the bias behavior of human observers implies that human estimation behavior can be understood by adaptive encoding and decoding processes.

### Time scales of the attractive and repulsive effects

We believe our results can provide a unified framework for interpreting the existing literature. For example, several studies have reported that the magnitude of attractive bias increases with increasing time delay between stimulus presentation and response (Fritsche et al., 2017; Bliss et al., 2017). These stronger attractive biases may be attributed to more uncertain sensory information corrupted by higher memory noise, while the magnitude of repulsion remains relatively the same, since the time delay took place after the sensory encoding and thus would have not affected the repulsive bias. In the extreme case where there is no time delay between the stimulus and response (and thus minimal memory noise), the net bias becomes repulsive (Bliss et al., 2017) as the magnitude of attraction becomes smaller than that of repulsion. Several studies have shown that estimation responses are attracted to immediately preceding stimuli and are repelled away from stimuli from many trials back (Fritsche et al., 2020; Gekas et al., 2019; Suárez-Pinilla et al., 2018). This time, both attraction and repulsion are likely to be affected by the timescale of the trial history. The relative speeds of decay may be each measured by increasing propagation noise with longer timescales (van Bergen & Jehee, 2019) and smaller gain reduction of sensory neurons by stimuli from many trials back compared to the immediately preceding one (Fritsche et al., 2021), respectively.

### Comparison to attraction and repulsion in non-dynamic stimuli

Our proposed model is comparable to the efficient encoding and optimal decoding theory for feature estimation under a static environmental distribution (Wei & Stocker, 2015). This theory proposes that the visual system efficiently encodes incoming information, allocating more neural resources to the representation of more probable stimulus values, which typically causes a bias away from the mean of the distribution. The system then decodes the sensory representations in an optimal manner by combining the representation with prior knowledge about the stimulus distribution, which typically results in a bias toward the mean of distribution. The relative magnitudes of these opposite effects determine the net bias in perceptual estimates. However, even in that work, there is no behavioral data that show both attractive and repulsive effects at the same time. A notable difference between the model for stationary environments and our proposed model for changing environments is that sensory adaptation for efficient coding, which is supposed to cause repulsive bias, is described as adapting to the preceding stimulus, as evidenced by the joint bias map and conditional bias plot (**Fig. 2a,c**). Consequently, it becomes possible to separately estimate the attractive bias, which is centered around the previous response, and repulsive bias, which is centered near the previous stimulus (**Fig. 4**).

### Sensitivity to temporal statistics

For the Bayesian observer to be truly optimal, it is necessary for the observer to maintain priors that approximate the true stimulus statistics of the external world (Ma, 2012). Can we learn novel temporal statistics that might differ from natural statistics and update priors accordingly? In this study, we divided subjects into groups with different propagation noise distributions: Some subjects encountered stimuli that were always very similar to the preceding one, while others encountered stimuli that were independent from the preceding one (see **Methods**). The main purpose was to increase the number of trials with dissociable stimulus and response directions, but our design also allowed us to test the effect of experimentally imposed temporal statistics on estimation behavior. If subjects learn new temporal statistics, then the magnitude of the biases would depend on those temporal statistics. The best-fitting parameters of the Stimulus & Response model showed that there was a significant effect of propagation noise level on the strength of repulsive bias away from the previous stimulus (*p* = 0.002; Kruskal-Wallis test), but no statistical significance was found in the attractive bias to the previous response (*p* = 0.283; **Supplementary Fig. 2**). The fitting results of the Attraction & Repulsion model showed similar patterns (*p* = 0.005 and *p* = 0.927, respectively). Indeed, priors specifying the statistical characteristics of the natural environment are often thought to be hard-wired through processes of evolution and development, and are difficult to update (Seriès & Seitz, 2013). In the case of temporal statistics, subjects can adapt their internal models of the temporal statistics to the stimulus set they have encountered, but still retain priors on positive temporal correlation within the stimulus set (Kwon & Knill, 2013). Future work is required to fully characterize whether and how priors on the temporal statistics of the natural environment can be overridden by experimentally imposed statistics.

In summary, we demonstrated that the perceived direction of motion in the current trial depends on both the perceived and presented direction of motion in the previous trial, but in opposite directions. Subjects’ estimates of the direction of motion were repelled away from stimuli that had recently been seen, while they were attracted toward responses that had recently been made. We found that the rich pattern of the sequential effects can be well described by the linear sum of two curves, each representing bias away from the sensory measurement of the previous stimulus and bias toward the previous response. This suggests that the mechanisms underlying the generation of attractive and repulsive sequential biases are largely independent of each other. Our findings suggest that the visual system adaptively encodes and decodes incoming sensory information by referring to the recent history of changing environments to optimize visual processing.

## Methods

### Subjects

Thirty-two subjects (16 females, aged 18 – 29 years) participated in the experiment. They were naïve to the purpose of the experiment and had not participated in similar experiments. We required subjects to have normal or corrected-to-normal vision and obtained written informed consent from all subjects prior to their participation. All procedures were approved by the Ulsan National Institute of Science and Technology Institutional Review Board.

### Procedure

**Figure 1a** illustrates the sequence of events in each trial. The subjects viewed the stimuli binocularly from a distance of 137 cm in a dark room, resting their head on a chinrest. Each trial began with the presentation of a fixation point. The subjects were instructed to fixate on the fixation point during the presentation of the motion stimulus. After 0.5 s, the motion stimulus was presented for another 0.5 s, followed by a 1.5-s delay during which only the fixation point was on the screen. After the delay, a circular ring appeared, and the subjects reported the perceived direction of motion by swiping a finger on a touchpad to extend a dark bar from the fixation point in the direction of motion that they had perceived and terminated the trial by clicking on the touchpad. Response time was not limited, and subjects made a response within 1.10 ± 0.05 s (mean ± s.e.m. across subjects). Trials were separated with a 1.5-s inter-trial interval during which the screen was blank.

The direction of motion of the current trial was solely determined with respect to the direction of motion of the immediately preceding trial. Specifically, subjects were divided into four groups of conditions in which the direction of stimulus motion was randomly varied from the direction of motion in the previous trial following uniform distribution with different ranges (±20°, 40°, 80°, or 180°). After one practice session, all subjects performed at least 1,500 trials over three sessions on consecutive days. Each session consisted of five blocks of roughly 100 trials each, lasting up to 60 minutes. After each block, the subjects received numerical feedback regarding their mean absolute response error.

### Stimuli

Stimuli were generated using MATLAB and the Psychophysics Toolbox (Brainard, 1997) and were displayed by a DLP projector (1920 × 1080; 120 Hz). All stimuli were presented at the center of a dark gray background of 20 cd/m^2^. The fixation point was a white circular point (diameter: 0.4°; luminance: 80 cd/m^2^). A gap of 1° between the fixation point and motion stimulus helped the subjects maintain fixation. The motion stimulus was a field of moving dots (diameter: 0.1°; luminance: 80 cd/m^2^) contained within a 5-degree circular aperture centered on the fixation point. The dots were plotted in three interleaved sets of equal size. Each set was plotted in one of three successive video frames and was shown for a single frame. Three frames later, randomly chosen 40% of dots from that set moved coherently in a designated direction at a speed of 4°/s; the remainder of the dots were replotted at random locations within the aperture. Dots that moved outside the aperture were placed on the opposite side of the aperture. Together, the three sets produced an average dot density of 48 dots/(deg^2^s). The presentation of a black circular ring (diameter: 6.6°; width: 0.15°; luminance: 15 cd/m^2^) centered on the fixation cued subjects to report their estimate, which they did by swiping their finger on a touchpad to extend and align a black bar (width: 0.15°) to the direction of their estimate and then clicking on the touchpad to confirm.

### Analysis

All analyses were performed using MATLAB and the CircStat Toolbox (Berens, 2009). In the first step of data analysis, we corrected for an individual subject’s idiosyncratic estimation bias to or away from the cardinal axes (Kosovicheva & Whitney, 2017; Wei & Stocker, 2015), which is unrelated to sequential effects, by fitting each subject’s response errors with a 10-degree polynomial function of stimulus direction (Manassi et al., 2018; van Bergen & Jehee, 2019). The residuals from this polynomial fit were used in the remaining analyses. We further excluded trials with response error more than 2.5 s.d. away from the subject’s mean response error and the subsequent trials on which subjects could have been affected by those preceding trials with outlying responses. In total, 4.02% of the trials were excluded, and the overall mean absolute response error was 6.99 ± 0.27° (mean ± s.e.m. across subjects).

The subjects’ dependencies on the direction of the previous stimulus and/or response in estimating the direction of the current stimulus were quantified by fitting the first derivative of a Gaussian (DoG) curve(s) to their response errors (Fischer & Whitney, 2014). The DoG curve is given by 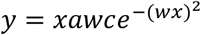 where *y* is the response error, *a* is the amplitude of the curve peaks, *w* scales the width of the curve, and *c* is a constant, 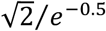. The input to the function, *x*, was either the relative direction of the previous stimulus (Stimulus model; **Fig. 4a**) or relative direction of the previous response (Response model; **Fig. 4b**). To effectively capture the characteristic pattern in the estimation data (**Fig. 2**), we fitted a linear sum of two independent DoG curves, one receiving as input the relative direction of the previous stimulus, and the other receiving the relative direction of the previous response (Stimulus & Response model; **Fig. 4c**). To explore the possibility that the centers of biases are not exactly located on the previous stimulus or response, we fitted a model similar to the Stimulus & Response model, but setting the inputs to the function, *x*_*att*_ and *x*_*rep*_, to be a combination of the previous stimulus and previous response, relative to the current stimulus (Attraction & Repulsion model). Specifically for the Attraction & Repulsion model, the bias amplitudes, *a*_*att*_ and *a*_*rep*_, were constrained to be positive and negative, respectively.

All parameters were estimated using a hierarchical Bayesian approach that uses the aggregated information from the entire population sample to inform and constrain the parameter estimates for each individual (Kruschke, 2014). Specifically, we assumed a hierarchical prior on parameters, in which the parameters for each subject were drawn from independent von Mises distributions characterizing the population distributions of the model parameters. Before being constrained by higher-level parameters, width parameters *w* were transformed to 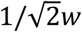, directly representing the peak location of the DoG curve, and concentration parameters *k* of the population distributions were transformed to 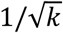, analogous to the standard deviation of the Gaussian distribution. Priors on the mean of the population distributions were set to broad uniform distributions with ranges large enough to cover all practically possible values. Priors on standard deviation of the population distributions were set to gamma distributions with parameters that made it vague on the scale of the data, as is common in the field (Kruschke, 2014).

We used a Markov chain Monte Carlo technique, specifically a Metropolis-Hastings algorithm, to compute the posterior probability density of the parameters. After using the first 10 million iterations as a burn-in period, we used the subsequent 10 million new samples from 10 independent chains to estimate the posterior probability density function. We further thinned the samples by selecting every 1,000 samples in the chain, resulting in a final set of 10,000 samples for each parameter and reducing autocorrelations in the samples to near zero. Convergence of the chains was confirmed by visual inspection of trace plots and Gelman-Rubin tests (Gelman & Rubin, 1992). All parameters in the model had 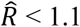, suggesting that all chains successfully converged to the target posterior distribution. For the statistical significance of the model parameters, we report as *p* values twice (i.e., two-tailed) the percentage of samples that have parameter values (or difference between parameter values) larger or smaller than zero. All *p* values in the main text, unless otherwise noted, were computed in this manner.

### Observer model

We begin with a conventional encoding model for a population of *N* = 20 sensory neurons responding to a stimulus direction *θ*. We assumed that the number of spikes emitted in a given time interval by the *i*th neuron is a sample from an independent Poisson process, with the mean rate determined by its tuning curve (von Mises distribution). Given these assumptions, the encoding model is specified as the probability of observing a particular population response **r** ≡ {*r*_*i*_, …, *r*_*N*_} for a given stimulus direction *θ*:

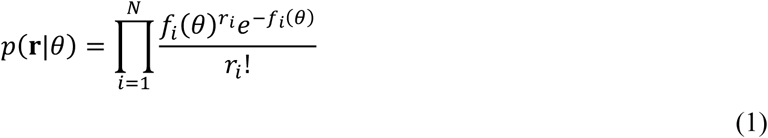

where *f*_*i*_(*θ*) = *g*_0_ exp(*κ* cos(*θ* – *θ*_*i*_)) is the tuning curve of the *i*th neuron with a response gain *g*_0_ = 4, a concentration parameter *κ* ≈ 3.65 (analogous to a Gaussian function with a 30° standard deviation), and the preferred direction *θ*_*i*_. Notably, from the perspective of the encoder that generates a noisy sensory response **r** in response to an unknown *θ*, this relationship becomes a function of *θ* with a fixed **r**, which is known as the likelihood function.

Sensory coding is not invariant to the temporal context, but is adaptive to the recently encountered stimulus. Based on neurophysiological studies, we assume that the primary effect of adaptation is a change in the response gain *g*_0_ of neuron *i*, such that those neurons most responsive to the adapter reduce their gain the most (Clifford, 2002; Clifford et al., 2007; Kohn, 2007; Webster, 2015). Specifically, we assume that the amount of gain reduction in the *i*th neuron is a von Mises function of the difference between the adapter direction and the preferred direction of that neuron (Seriès et al., 2009)

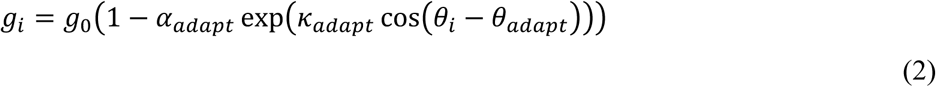

where *α*_*adapt*_ is an adaptation ratio specifying maximal suppression, and *κ*_*adapt*_ is a concentration parameter that determines the spatial extent of response suppression in the direction domain. For the model simulation, the gain reduction due to adaptation was modulated by *α*_*adapt*_ = 0.35 and *κ*_*adapt*_ = 8.2. The encoding model after adaptation is illustrated in **Figure 6a** and **b**.

One concern that can be raised here is that sensory neurons should adapt to the sensory measurement rather than the physical stimulus because the neurons would not have direct access to the true stimulus value. To this end, we also tested an alternative mechanism in which the response gain of a neuron was reduced according to the number of spikes emitted by the neuron in response to the adaptor:

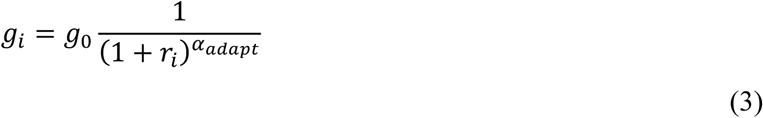

where the role of *α*_*adapt*_ is qualitatively similar to that in equation (2). We confirmed that changing the adaptation scheme from equation (2) to equation (3) does not change the simulation results: the average likelihood function is shifted away from the adaptor direction.

Next, we proceed with the notion that the natural environment is stable over time, and the model observer incorporates such temporal statistics by performing recursive Bayesian inference (**Fig. 6c**). Specifically, the model observer knows that in the natural environment, the stimulus variable *θ* propagates from *θ*_*t*−1_ to *θ*_*t*_ in a given time interval, following a probability distribution called the propagation noise distribution. Motivated by previous work (van Bergen & Jehee, 2019), the propagation noise was modeled as a weighted average of von Mises and uniform distributions

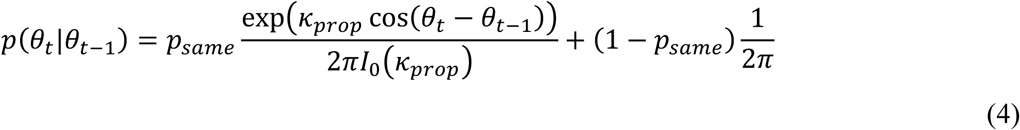

where *I*_0_(·) is the modified Bessel function of the order of 0. We set *p*_*same*_ = 0.9 and *κ*_*prop*_ = 14.6, such that the resulting distribution closely approximates the empirical observations (van Bergen & Jehee, 2019). For an observer who has already inferred the previous stimulus and knows that the stimulus value would change following the propagation noise distribution, it is reasonable to predict the current stimulus by convolving the distribution of knowledge about the previous stimulus, *p*(*θ*_*t*−1_|**r**_1: *t*−1_), with the distribution of changes that can occur between consecutive time points, *p*(*θ*_*t*_|*θ*_*t*−1_), as follows:

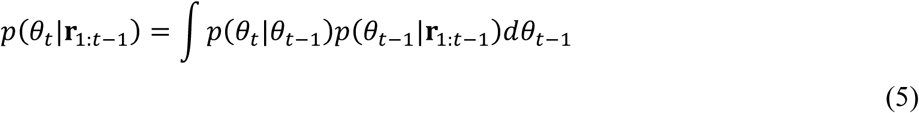

Finally, to infer the (unknown) stimulus value *θ*_*t*_, the observer combines prediction from the past sensory responses (equation (5) with the current sensory likelihood (equation (1), according to Bayes’ rule:

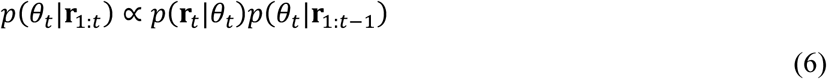

This final distribution characterizes the model observer’s posterior belief on the current stimulus, based on all the information available to the model observer at time *t*, including information obtained from both past and current sensory responses. We assume a squared-error loss function (or *L*_2_ norm), which is equivalent to computing the posterior mean (Jazayeri & Shadlen, 2010).

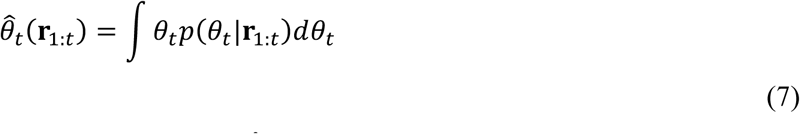

The model observer’s final estimate of the stimulus direction, 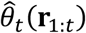, is a function of sensory response **r** over all past and current trials, which makes the marginalization over the latent variable **r** particularly demanding. Specifically, to compute the distribution of estimates 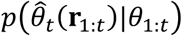 on a given trial, one would have to marginalize over all possible combinations of **r**_1:*t*_ in response to a stimulus sequence *θ*_1:*t*_, which grows exponentially with the number of trials. Therefore, we simulated the estimation behavior of the model observer using parameters fitted by hand to human data and demonstrate that the model can generate the characteristic pattern inherent in human estimation data. Direction estimates of the model observer were obtained using the same sequence of stimuli that our subjects encountered, assuming no motor noise. We analyzed the simulation data as we did with the empirical data, except for the cardinal bias correction. The results are shown in **Figure 6d**.

## Acknowledgements

This research was supported by the National Research Foundation of Korea (NRF-2020S1A3A2A02097375 to O.-S.K.).

