## Supplementary Figure 1, Supplementary Fig. 2 for "Dissecting the effects of adaptive encoding and predictive inference on a single perceptual estimation"

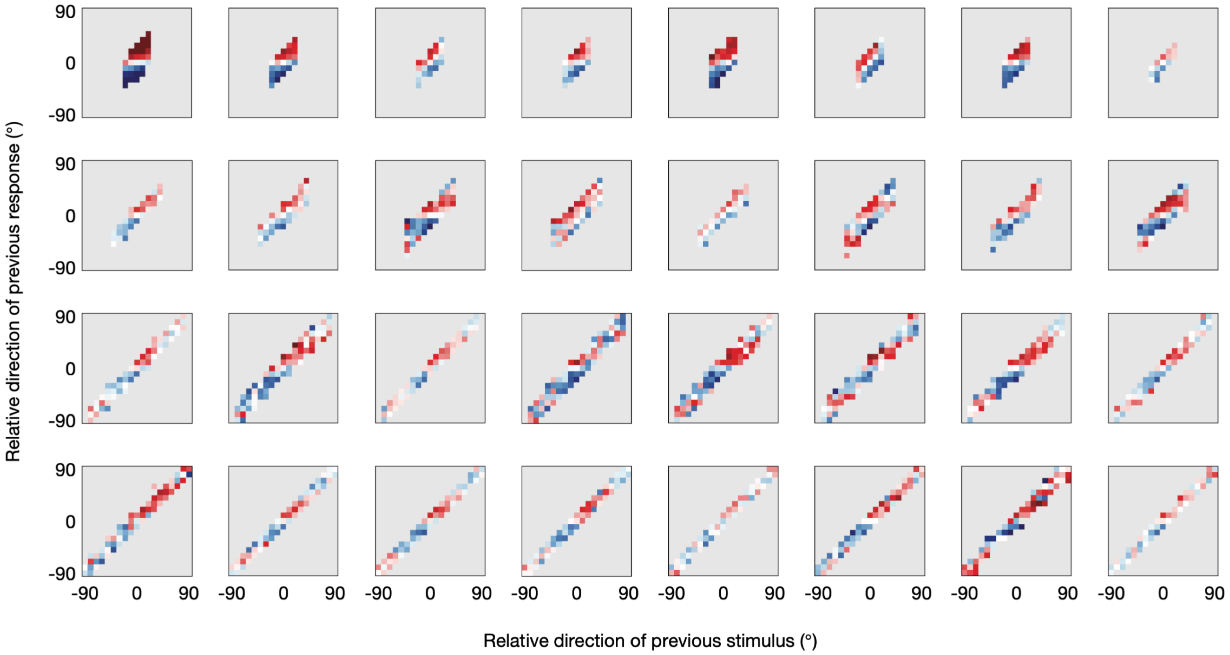


**Supplementary Figure 1. Joint bias map for individual subjects.** Color conventions are as described in **Figure 2a**, and gray regions indicate data not available.

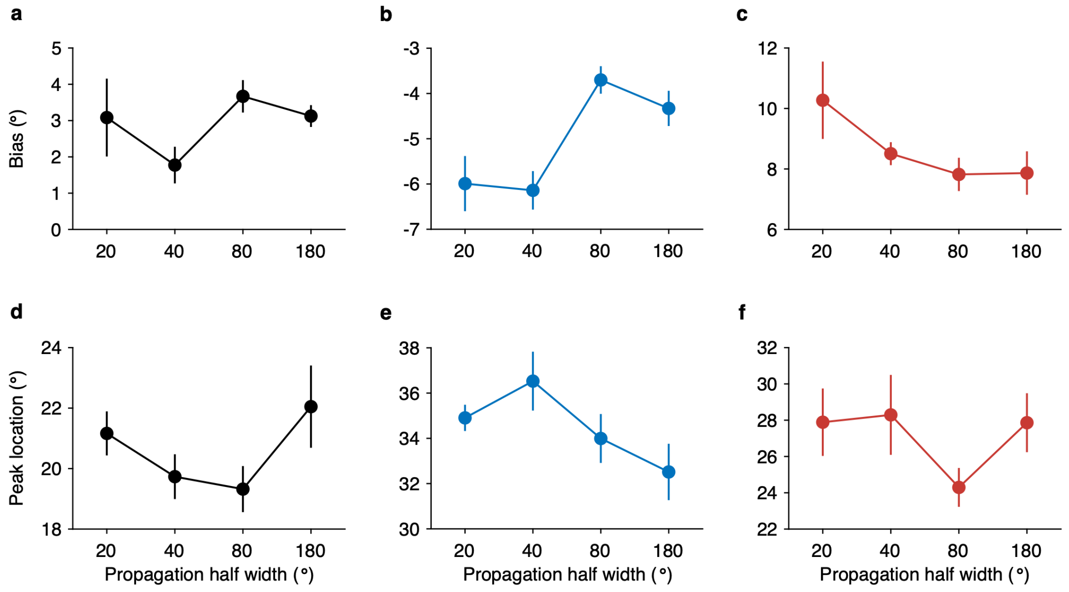


**Supplementary Figure 2. Minimal group difference in model parameters. (a)** Group difference in bias magnitudes estimated by the Stimulus model. There was no significant effect of propagation noise ($p$ = 0.148; Kruskal-Wallis test). **(b)** Group difference in magnitudes of bias away from the previous stimulus, estimated by the stimulus & Response model. There was a significant main effect of propagation noise on the bias magnitude ($p$ = 0.002). A post hoc Tukey test showed that only the 80° group had significantly stronger repulsive biases compared to the 20° group ($p$ = 0.016) and the 40° group ($p$ = 0.005). **(c)** Group difference in magnitudes of bias toward the previous response, estimated by the Stimulus & Response model. Although the bias magnitudes qualitatively decreased with increasing propagation noise, no statistical significance was found ($p$ = 0.283). Error bars represent 95% confidence intervals. **(d-f)** Same as in **a-c** but for group differences in locations of maximum biases. No statistical significance was found in peak locations estimated by the Stimulus model ($p$ = 0.232) and peak locations of repulsive biases ($p$ = 0.268) and of attractive biases ($p$ = 0.378) estimated by the Stimulus & Response model.
